# Control of artefactual variation in reported inter-sample relatedness during clinical use of a *Mycobacterium tuberculosis* sequencing pipeline

**DOI:** 10.1101/252460

**Authors:** David H Wyllie, Nicholas Sanderson, Richard Myers, Tim Peto, Esther Robinson, Derrick W Crook, Grace Smith, A Sarah Walker

**Affiliations:** Nuffield Department of Medicine, John Radcliffe Hospital, Headley Way, Oxford OX3 9DU, UK; Public Health England Academic Collaborating Centre, John Radcliffe Hospital, Headley Way, Oxford OX3 9DU, UK; The National Institute for Health Research Health Protection Research Unit (NIHR HPRU) in Healthcare Associated Infections and Antimicrobial Resistance at University of Oxford; Department of Bioinformatics, Public Health England, 61 Colindale Avenue, London NW9 5EQ, UK; Public Health England National Regional Mycobacteriology Laboratory North and Midlands, Heartlands Hospital, Birmingham BS9 5SS

## Abstract

Contact tracing requires reliable identification of closely related bacterial isolates. When we noticed the reporting of artefactual variation between *M. tuberculosis* isolates during routine next generation sequencing of *Mycobacterium spp,* we investigated its basis in 2,018 consecutive *M. tuberculosis* isolates. In the routine process used, clinical samples were decontaminated and inoculated into broth cultures; from positive broth cultures DNA was extracted, sequenced, reads mapped, and consensus sequences determined. We investigated the process of consensus sequence determination, which selects the most common nucleotide at each position. Having determined the high-quality read depth and depth of minor variants across 8,006 *M. tuberculosis* genomic regions, we quantified the relationship between the minor variant depth and the amount of non-Mycobacterial bacterial DNA, which originates from commensal microbes killed during sample decontamination. In the presence of non-Mycobacterial bacterial DNA, we found significant increases in minor variant frequencies of more than 1.5 fold in 242 regions covering 5.1% of the *M. tuberculosis* genome. Included within these were four high variation regions strongly influenced by the amount of non-Mycobacterial bacterial DNA. Excluding these four regions from pairwise distance comparisons reduced biologically implausible variation from 5.2% to 0% in an independent validation set derived from 226 individuals. Thus, we have demonstrated an approach identifying critical genomic regions contributing to clinically relevant artefactual variation in bacterial similarity searches. The approach described monitors the outputs of the complex multi-step laboratory and bioinformatics process, allows periodic process adjustments, and will have application to quality control of routine bacterial genomics.

## INTRODUCTION

Identifying closely related bacterial isolates is required for clinical and epidemiological purposes [1–3]. Most published approaches using short-read next generation sequencing (NGS) rely on mapping to a high-quality reference sequence followed by consensus base calling [1–8]. A known problem with this approach concerns the existence in many bacterial genomes of ‘hard-to-map’ regions which are either repeated within the genome, or which contain regions of low sequence complexity. High-confidence mapping of short reads to such regions is difficult or impossible, and so determining the consensus sequence of these regions is difficult. One approach to managing this problem is to identify these regions bioinformatically prior to mapping by analysis of sequence complexity [9], or from repetitiveness within the genome [10]. Base calls within these pre-specified regions are then ignored (‘masking’) when assessing relatedness of isolate sequences. A second complementary approach filters base calls based on read mapping confidence as reported by various mappers [11–14] in the form of MAQ scores.

*Mycobacterium tuberculosis* is one of the most important pathogens of humans, with about 3 million cases of TB confirmed by culture globally each year [15]. Recently, laboratory protocols have been described and deployed by Public Health England [4] in which the species and drug resistance of *Mycobacteria,* including *M. tuberculosis*, are identified by sequencing microbial DNA. Laboratory processing of clinical samples suspected of containing *Mycobacteria* involves decontamination using chemicals which kill non-Mycobacterial species before the samples are inoculated into broth culture [16]. Mycobacterial Growth Indicator Tubes (MGIT) and associated tube monitoring equipment are a commercially available implementation of such a broth culture system.

In the process adopted by Public Health England, sequencing and bioinformatics analysis of DNA extracted from positive MGIT tubes allows determination of Mycobacterial species and drug resistance [17]. This laboratory and bioinformatic process also allows the genetic distance between *M. tuberculosis* isolates to be estimated, using sequences derived by consensus base calling from mapped data. The organism co-evolved with human populations as they migrated, generating multiple lineages which differ from the ancestral sequence by hundreds or thousands of single nucleotide polymorphisms [18], as well as small indels, gene deletions and inversions [19]. However, the evolutionary clock rate of the organism is slow at about 0.5 single nucleotide variants (SNV)/genome/annum [3, 5, 7] and small numbers of single nucleotide variants (SNV) are of clinical significance: studies based on retrospective collections of *M. tuberculosis,* grown on solid media prior to sequencing, have proposed thresholds of 5 SNVs as compatible with recent transmission [3, 5, 7]. The bioinformatic processes used for relatedness estimation in the deployed pipeline were also optimised using samples re-grown from frozen stocks on solid media (Lowenstein-Jensen slopes) [16].

Quality of complex processes deployed in medical laboratories is assured by adherence to quality standards, such as those laid out in ISO15189 [20]. These standards require that the processes followed, and the environments in which they operate, comply with to patterns of work known to enhance the consistency and interpretability of the laboratory outputs. For example, in a drug testing laboratory a set of samples of known composition may be run through the analysers to confirm that particular commonly found substances which might potentially interfere with the assay (such as caffeine, paracetamol, and so on) have no impact on detection of the drug of interest. *M. tuberculosis* infection is commonly diagnosed from sputum samples, which contains a wide variety of organisms other than *Mycobacteria* [21]. DNA from such organisms may contain sequences homologous with those present in *Mycobacteria*, for example in highly conserved core bacterial genes. As such, this non-Mycobacterial DNA has the potential to interfere with assays based on mapping of Mycobacterial reference genome mapping.

Here, we investigate the concept of interfering substances in the context of the detection of closely related *M. tuberculosis* isolates. To do so, we describe a process which we call *adaptive masking.* This defines ‘hard-to-map’ regions by adapting the selection of such regions to the complex laboratory, sequencing and consensus base calling process, which operates independently of both *ab initio* sequence examination and filtering based on reported mapping quality. Our work was motivated in part by observations from analysis of prospective sequencing of *M. tuberculosis* sequences in England using a previously described bioinformatics pipeline [17]. It appeared that high SNV distances were being reported between isolates with a strong epidemiological likelihood of having recently transmitted to each (e.g. isolates with unusual resistance profiles from individuals who were co-habiting): that is, false positive variation between isolates was being reported. We assess the impact of adaptive masking on addressing this problem, and discuss quality control of relatedness monitoring in the context of continuous process monitoring in accredited clinical laboratories.

## METHODS

### Isolation of DNA from Mycobacteria and Sequencing

Clinical specimens were decontaminated and inoculated into Mycobacterial Growth Indicator Tubes (MGIT) tubes [16]. Positive samples were extracted [4]. After DNA extraction, Illumina sequencing libraries were prepared using either 11 (early in the study) or 15 (later in the study) Mycobacterial DNA extracts, as previously described [4]. All samples sent from patients for processing for *Mycobacteria* between 1/05/2016 and 30/05/2017 to a single reference laboratory were studied; the catchment of this laboratory is approximately 15 million people, or about one third of England.

### Bioinformatic processing

Reads obtained from the MiSeq instrument were first examined for the presence of *Mycobacterium tuberculosis* using the Mykrobe tool which detects species-specific k-mers [22]. Additional read classification was performed with Kraken [23], which assigns reads to the phylogenic tree of life. The database used was that described [24], with k-mer reduction to 25 gigabases. The total number of reads from bacterial genera other than *Mycobacterium* were determined [23], and human reads counted and then discarded.

Reads were mapped to the H37Rv v2 genome (Genbank NC_000962.2) using Stampy [14], as described [4]. Samtools [25] was used to assess sequencing and mapping quality: high quality bases were considered to be those passing the –q30 and–Q30 30 filters (read quality and mapping quality all >30). Consensus sequence was called, requiring a minimum-read depth of 5, including at least one read on each strand. Where an alternative base represented more than 10% of read depth, the base was recorded as uncertain, as described [26].

The variant call format (VCF) file was parsed with custom python scripts, and the number of high quality bases (defined using the filters above) at each position determined. These frequencies were extracted, stored and indexed using SQLite using a python API constructed to allow extraction of mixture frequencies in arbitrary positions (https://github.com/davidhwyllie/BUGMIX). For selected regions, the proportion of mapped reads assigned to the *M. tuberculosis* taxon (NCBI taxid: 77643) by Kraken [23] was determined.

### Modelling minor variant frequencies

We determined the most common (major) variant at each position. All other variants are considered minor variants. We define

*n* - the total sequencing depth at one base;

*m* - the depth of most common variant at one base;

*m’* - the depth of all variants other than most common, *n*-*m* (Suppl. Fig. S1).

We divided up the H37Rv genome based on the annotation in NC_000962.3, identifying 8007 regions R, comprising open reading frames and regions between open reading frames (Supplementary Data S1).

For each of these regions *j*=1‥8007, if the region has length *l_j_*, the total number of minor variant bases V_j_ across the i=1‥l*_j_* bases in the region is given by equation (1) and the total read depth D_j_ across the gene by equation (2).

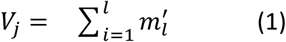

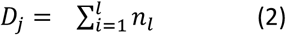

In order to describe the relationship between non-Mycobacterial DNA quantifications and minor allele frequency, we stratified the number of reads from each sample identified as being from bacterial genera other than *Mycobacterium (b)* into four approximately equally sized strata:

*b* < 1%,

1% ≤*b* < 5%,

5%*b* < 20%,

*b* ≥ 20%.

We constructed separate Poisson regression models relating minor base counts (V) for each of the 8,007 regions (with log link and offset log(D)) to the non-Mycobacterial bacterial read categories *b* (reference category < 1%), excluding any samples with zero high-quality depth in that region. We applied Bonferroni correction to model outputs to control for multiple testing ( α = 0.01/8007 = 1.2x10^−6^).

### Comparing impact of masking on pairwise comparisons

Based on analysis of model output (see Results), regions with higher minor variant counts than expected were identified. These regions were excluded from pairwise comparisons performed using the findNeighbour2 tool [27].

### Ethical framework

Public health action taken as a result of notification and surveillance is one of the Public Health England's key roles as stated in the Health and Social Care Act 2012 and subsequent Government directives which provide the mandate and legislative basis to undertake necessary follow-up. Part of this follow-up is identification of epidemiological and molecular links between cases. This work is part of service development carried out under this framework, and as such explicit ethical approval is unnecessary.

## RESULTS

### Samples studied

The PHE National Mycobacteriology Reference Laboratory Midlands implemented a laboratory process in which specimens received are decontaminated, and inoculated in MGIT bottles, DNA extracts from positive MGIT bottles made, and their contents determined using Illumina short read sequencing [17]. Using this process, in the 13 months from 1 May 2016 to 30 May 2017, *M. tuberculosis* was identified in 2,751 samples, sent from 2,252 patients (Fig. 1). Using these samples we derived and validated using an independent validation set (Fig.1) a strategy for investigating and controlling false positive variation between samples, which we here term *adaptive masking* (Fig. 2). The initial stages of adaptive masking involve estimating minor variant frequencies across the genome from mapped data, and determining whether these are related to the amount of non-Mycobacterial DNA present.

**Figure 1.**
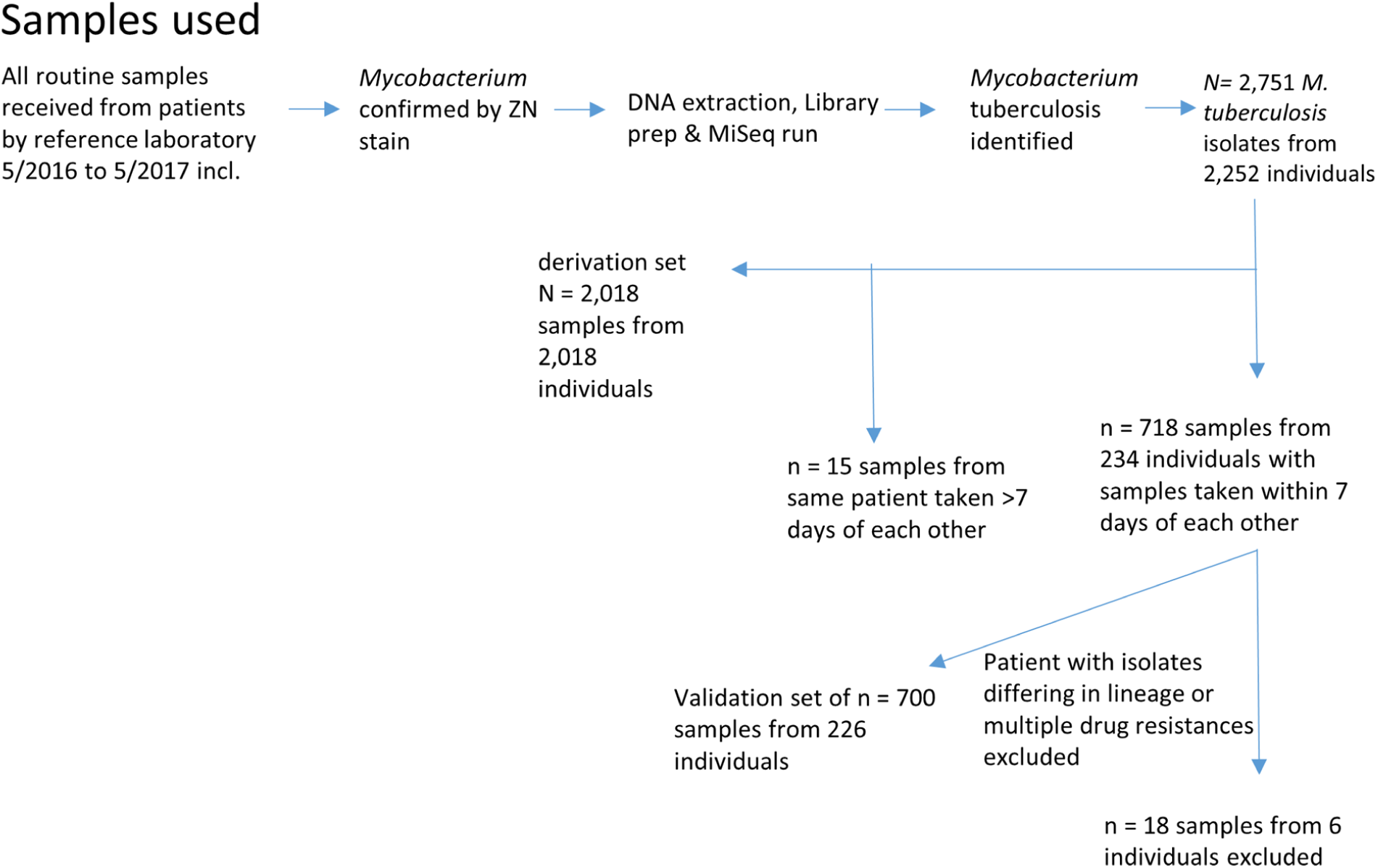
Samples used, derivation and validation sets. Legend: A flow chart describing the samples used, and the selection of derivation and validation sets.

**Figure 2.**
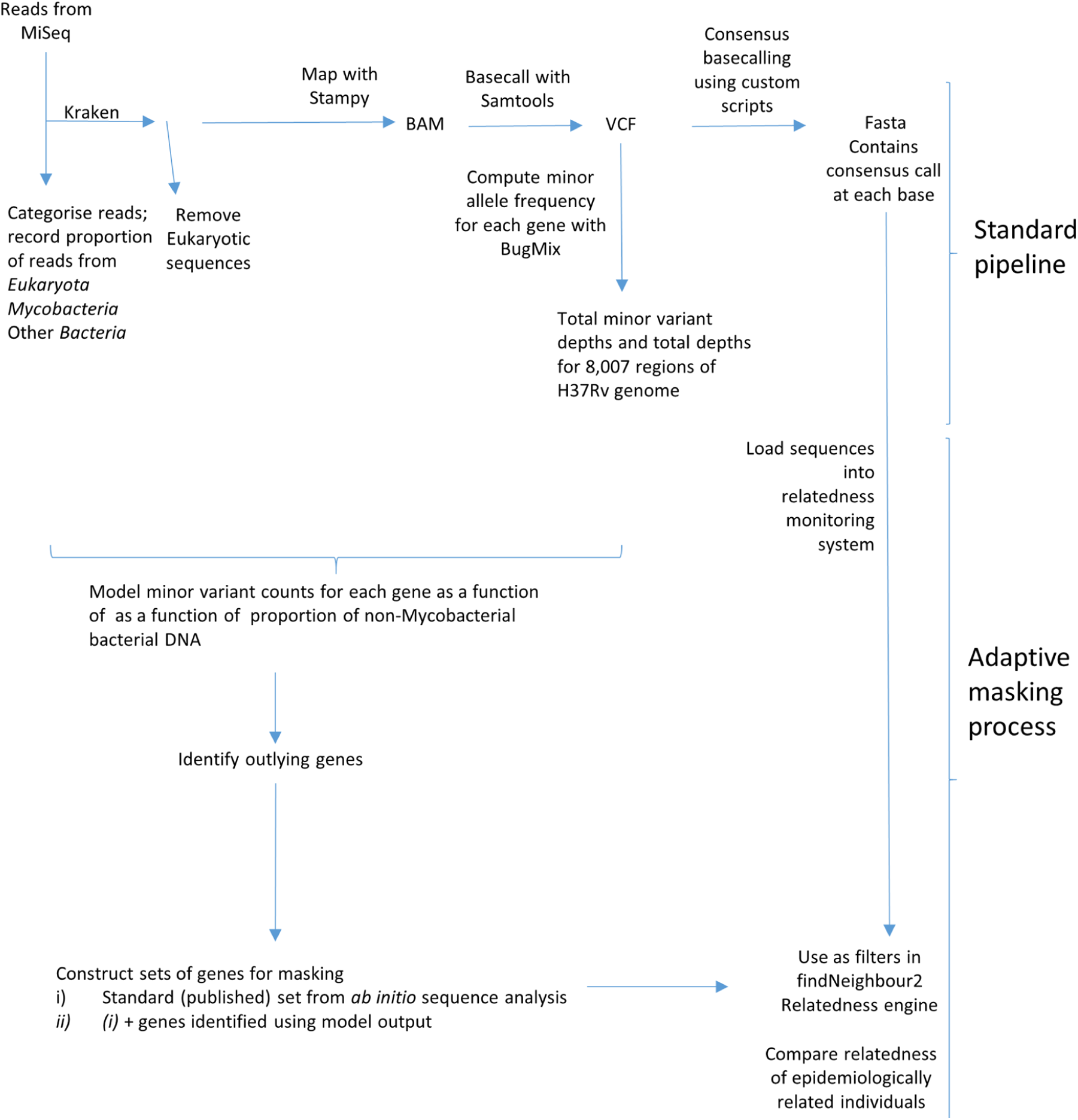
Bioinformatic Processes. Legend: A flow diagram illustrating the standard bioinformatics pipeline used, as well as the adaptive masking process used.

We identified 718 samples from 234 individuals from whom more than one positive sample had been obtained with 7 days of another. Of these, in six individuals samples were either reported as being of different lineages, as defined [28], differed in multiple drug resistances, or differed by >400 high quality SNVs. These observations we considered likely due to laboratory or sampling mix-ups, and samples from these patients were excluded. The other 700 samples were used as an independent validation set. From remaining samples, we identified the first sample from each of 2,018 individuals which were used to develop the adaptive masking strategy (Fig. 1).

### Quantifying extraneous DNA and minor variant frequencies post mapping

We determined the proportion of non-Mycobacterial bacterial DNA in each sample using Kraken [23], mapped all reads to the H37Rv reference genome irrespective of Kraken results, and filtered the mapped data using stringent quality filters such that the expected error rate is expected to be less than 10^−3^ (see Methods). We defined 8,007 genomic regions in the reference genome; these regions comprise all canonical open reading frames and the genomic regions between them (Supplementary Data S1). We were unable to assess one 15nt region between two PPE family members (positions: 3380439‥3380453) as no high quality data mapped there in any sample.

In the other 8,006 regions, we observed that both minor variant frequencies and the relationship between minor variant frequency and the number of reads of non-Mycobacterial origin differed markedly by gene. For example, the *B55* and *esxW* genes had respectively very low and very high minor variant frequencies, independent of non-Mycobacterial DNA quantity. A small group of genes, of which the ribosomal component *rrs* is an example, showed low minor variant frequencies, except when non-Mycobacterial DNA was present (Figure 3A, B).

**Figure 3.**
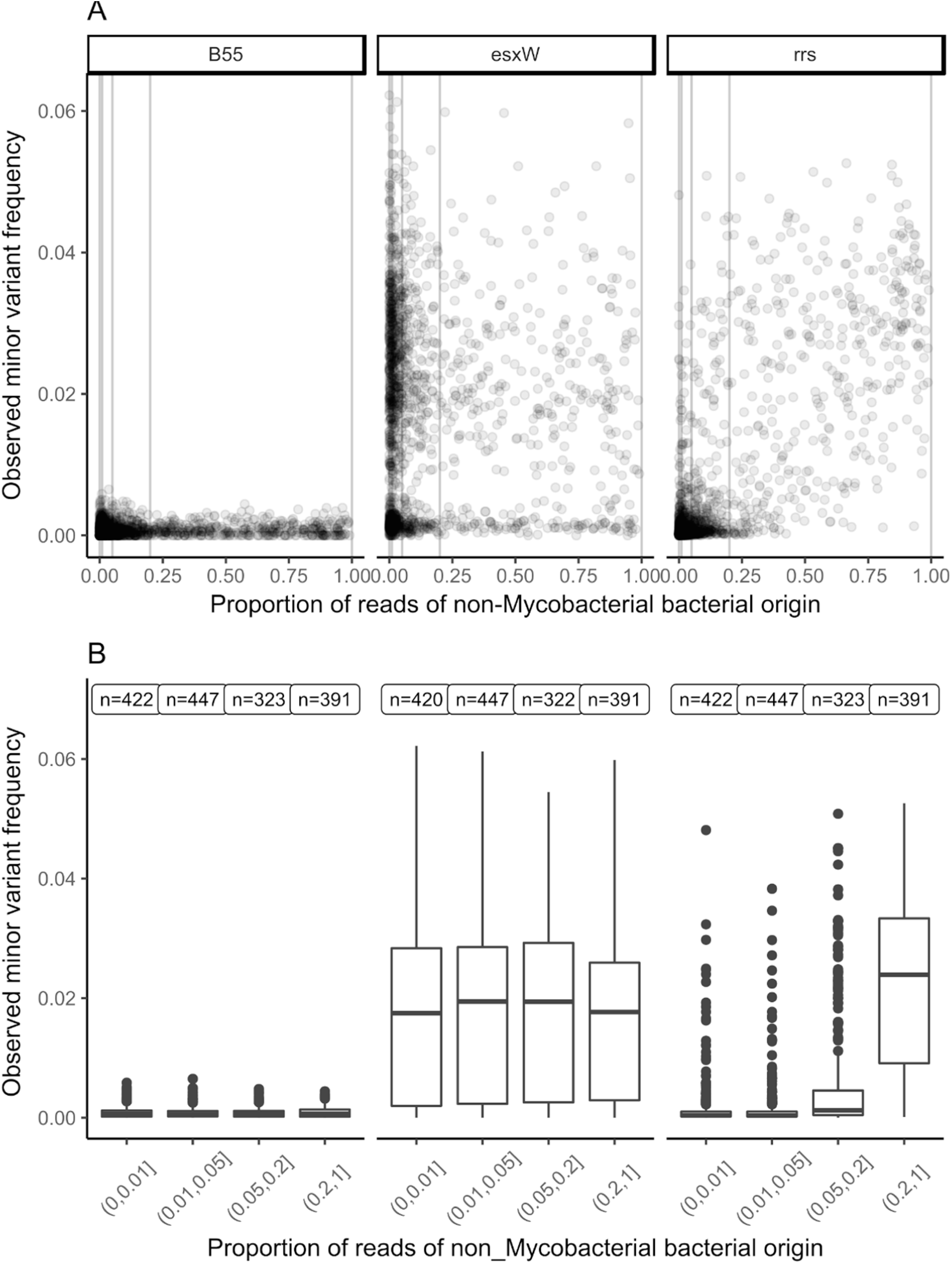
Minor variant frequency and non-Mycobacterial bacterial DNA quantities. Legend: The observed minor variant frequency for three regions of the *M. tuberculosis* genome (genes *B55*, *eswX*, and *rrs*) *vs*. the proportion of reads of non-Mycobacterial bacterial origin (as determined by Kraken) is shown for samples in the derivation set (n=2018). Panel (A) shows a dot plot, whereas in panel (B) the proportion of reads of non-Mycobacterial bacterial origin is stratified into strata with 1%, 5% and 20% boundaries. Numbers above each stratum refer to the number of samples with non-zero read depth in that region.

### Estimating the impact of extraneous bacterial DNA

We modelled the relationship between minor variant counts and the number of non-Mycobacterial reads, divided into four approximately equal sized strata (Figure 3B), using Poisson models (Suppl. Data S2). Separate models were constructed for each region. Estimated minor variant frequencies in samples with <1% non-Mycobacterial bacterial reads had a median of 5x10^−4^ (Fig. 4A) across the 8006 genomic regions, which is compatible with the expected mapping error rate of <10^−3^, given the filters applied.

**Figure 4.**
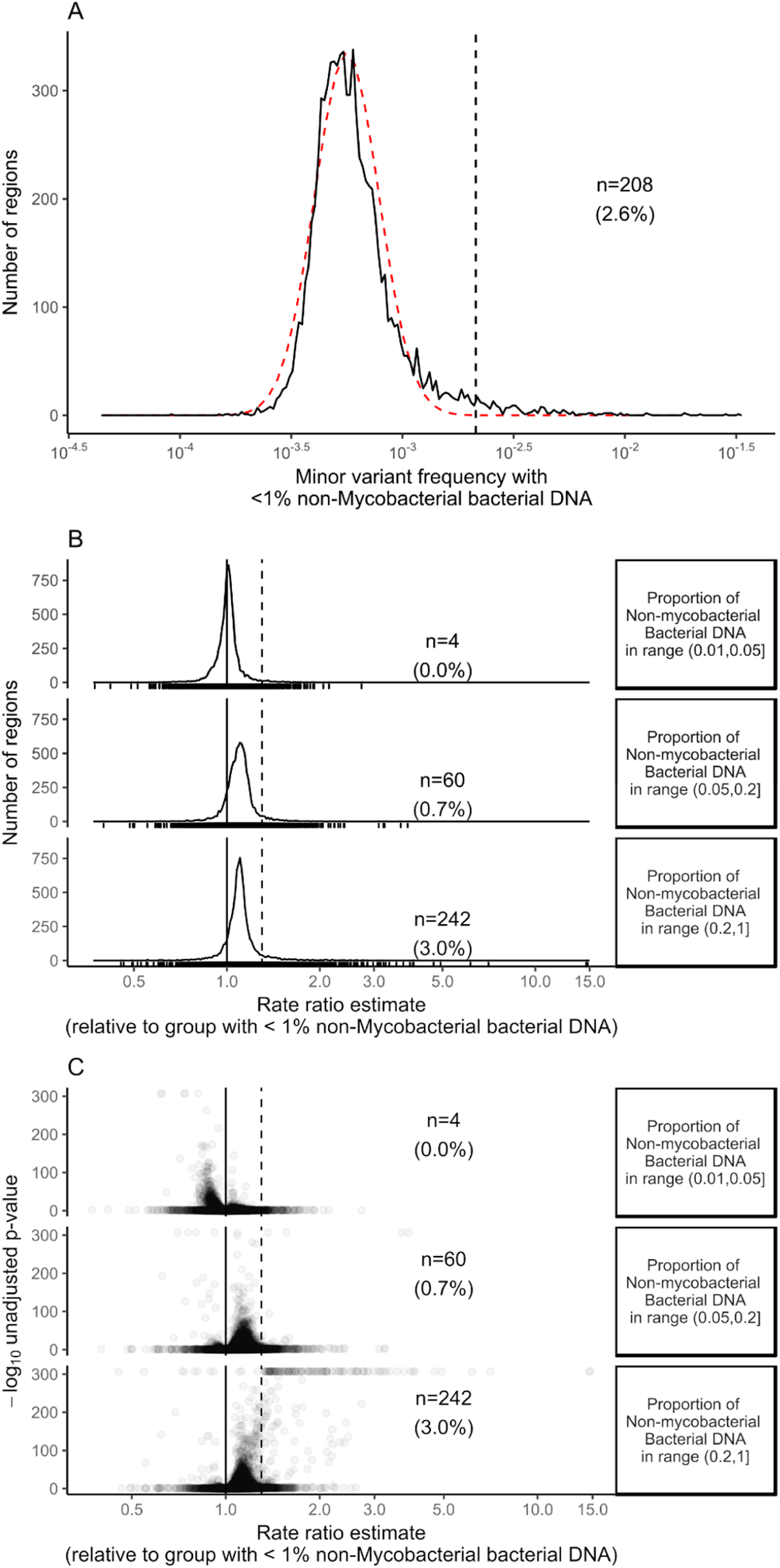
Modelling minor variant frequencies. Legend: For 8,006 genomic regions of the H37Rv reference genome, Poisson models were used to estimate the mean minor variant frequency. The estimated minor variant frequency when less than 1% non-Mycobacterial bacterial DNA is present (n=208, 2.6% regions) is shown in (A). The red line is a log normal distribution with μ = log(minor variant frequency with &1% non-Mycobacterial DNA) and σ = median absolute deviation(log(minor variant frequency with &1% non-Mycobacterial DNA)). In (B) the rate ratio estimates (i.e. the fold change associated with increases in non-Mycobacterial bacterial DNA quantifications) for each gene are shown. (C) shows the significance of a test comparing the log(rate ratio estimates) with zero, in the form of a Volcano plot. The dotted lines in (B) and (C) correspond to a 50% increase in rate ratio

The distribution of minor variant frequencies when less than 1% non-Mycobacterial bacterial DNA was present approximated a log-normal distribution with mean log(5x10^−4^) and standard deviation equal to the median absolute deviation (Fig. 4A, observed: black line; fitted, red line), but with a tail to the right. 208 regions (2.6% of the total 8006 regions), including esxW as well as other esx and PPE family members, had estimated minor variant frequencies > 2.1 x 10^−3^ when <1% non- Mycobacterial bacterial reads were present (Fig. 4A, Suppl. Data S2). This cutoff represents four median absolute deviations above the median; if the data were log-normally distributed, 24 samples would be expected with minor variant frequencies greater than this, vs. the 208 observed.

Overall, estimated minor variant counts rose as non-Mycobacterial DNA concentration rose, but for most regions the increase was small (Fig 4B): the median fold change in minor variant counts in the presence of >20% non-Mycobacterial DNA vs. <1% non-Mycobacterial DNA was 1.097 (i.e. a 9.7% increase, interquartile range 5.4% to 14.0%). 242 regions (3.0%) had statistically significant increases (Fig. 4B, 4C) of more than 50%. Most of these regions had the highest minor variant counts when >20% non-Mycobacterial DNA was present, although a small number had similar minor variant counts in the 5-20% range and the >20% range (Suppl. Fig. S2B).

### Mutually exclusive regions with increased minor variant frequency

Comparing regions with increased minor variant rates with low (<1%) non-Mycobacterial bacterial DNA with those with increased minor variant rates with high (>20%) non-Mycobacterial bacterial DNA shows these regions to be mutually exclusive (Fig. 5). The former include PPE and *esx* family members, while the latter include ribosomal components (*rrl*, *rrs*, *rplB*, and *rps* genes) as well as other highly conserved bacterial genes (tRNA genes, *fusA1*, *infA*, *dnaK*, and others) (Fig. 5, Suppl. Data S2). Examining Kraken read assignments in these highly conserved genes indicated that many reads mapping to these highly conserved regions cannot be unambiguously assigned to the *M. tuberculosis* taxon (Supp. Fig. 3), compatible with their high conservation across the tree of life.

**Figure 5.**
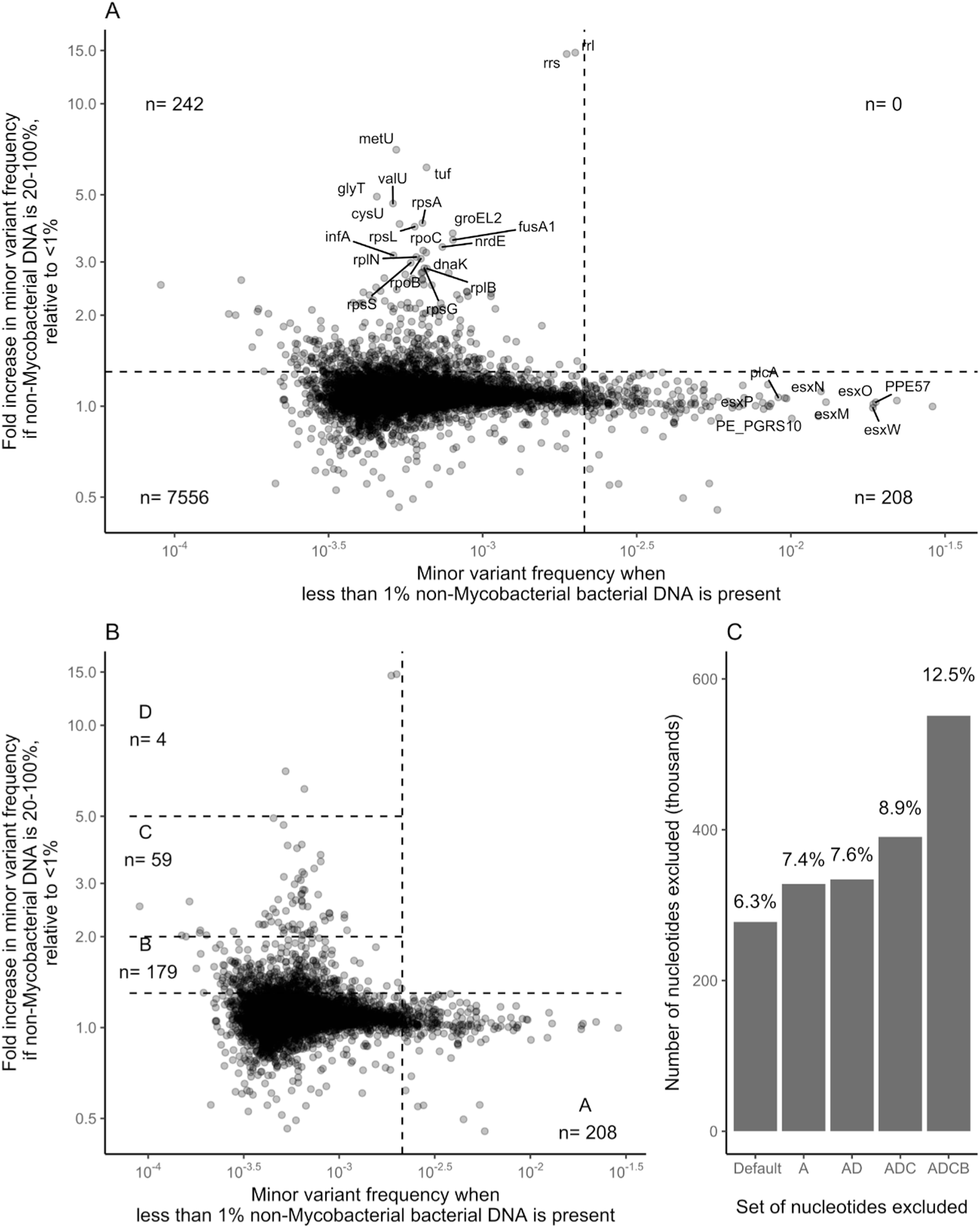
A distinct subset of genes are impacted by quantity of non-Mycobacterial DNA. Legend: (A) Fold change in minor variant frequency with & 20% non-Mycobacterial bacterial DNA present vs. &1% non-Mycobacterial bacterial DNA. Quadrant boundary markers correspond to (horizontal line) a 50% increase over &1% non-Mycobacterial bacterial DNA and (vertical line) a minor variant frequency of 2.1x10^−3^. (B) Genes with elevated minor variant frequencies when non-Mycobacterial bacterial DNA is low (<1%) or high (>20%) fall into mutually exclusive sets. (C) the number of bases represented by the deployed masking, vs. the deployed masking plus the genes in regions A, A+D, A+D+C, and A+D+C+B.

### Adaptive masking reduces the reporting of biologically implausible inter-individual variation

A published strategy for excluding regions of high mapping variation within the *M. tuberculosis* genome strategy masks (i.e. excludes from relatedness computations) 277,709 nt (6.3%) of the genome [4]. Excluding regions with high estimated minor variant counts with <1% non-Mycobacterial DNA (region A in Fig. 5B) adds an additional 1.1%. Excluding regions with increased estimated minor variant counts only in the presence of >20% non-Mycobacterial bacterial DNA (regions B-D) masks between 0.2% and 5.1% extra (Fig 5B,C). The masking of regions identified by ‘adapting’ to variation generated during the process forms the final part of the adaptive masking process.

In a validation set comprising isolates taken with 7 days of each other from 234 individuals, using the published strategy, 18/346 (5.2%) of pairs studied had ≥ 5 SNV variation, of which 10/346 were ≥ 20SNV. On exclusion of region D, which comprises the four genes most influenced by non-Mycobacterial DNA, all encoding ribosome associated products (the genes *rrl* and *rrs*, together with the tRNA *metU*, and the highly conserved bacterial gene *tuf*), 0/346 pairs differed by >= 5 SNP (p < 10^−4^ compared with the published method, Wilcoxon test on pairs). Additional exclusion of genes in regions B and C, mapping to which is less influenced by non-Mycobacterial DNA, had a limited impact (Figure 6).

**Figure 6.**
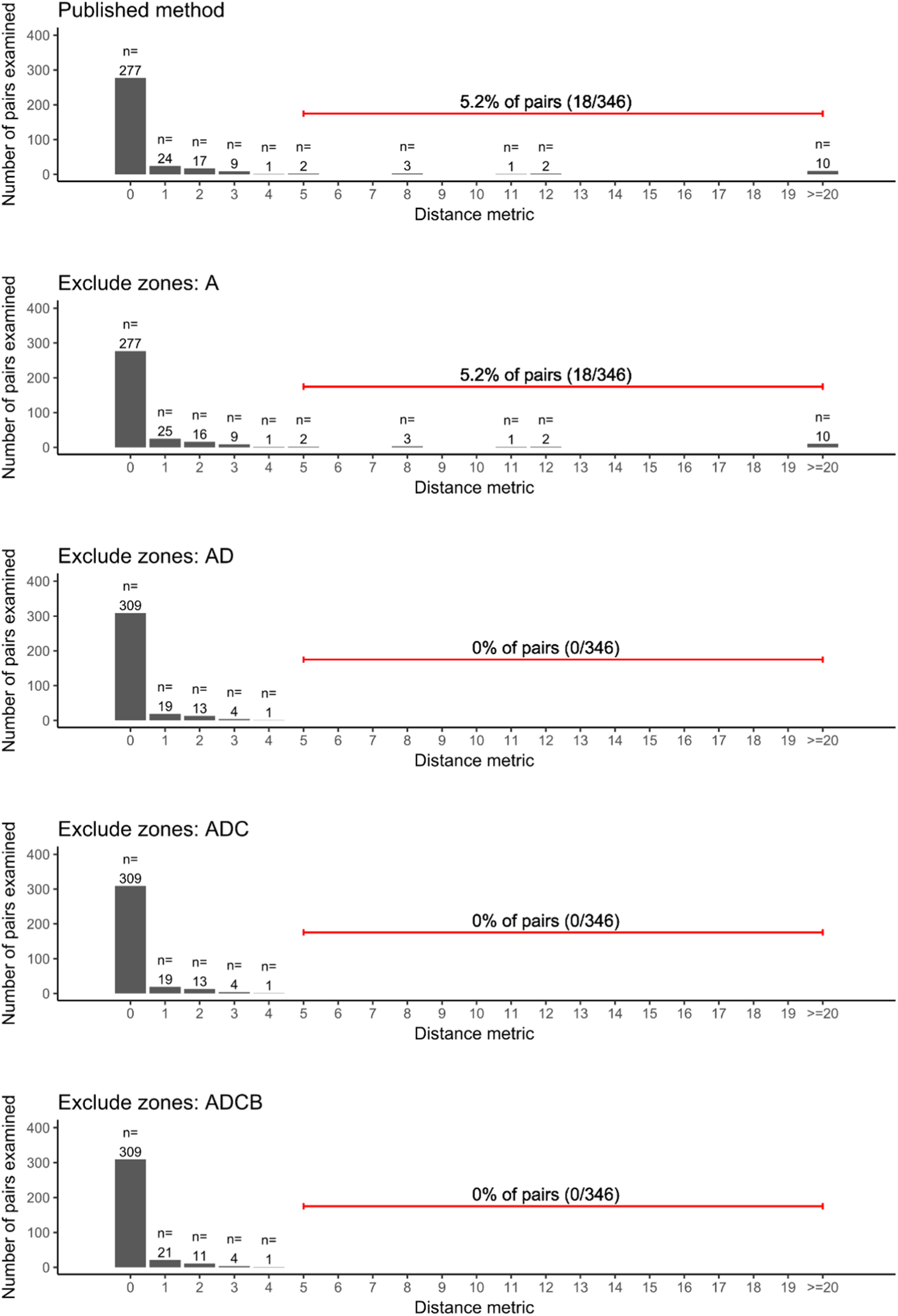
Impact of masking strategies on reported distances between closely related samples. Legend: SNV distances between pairs of *M. tuberculosis* genomes isolates from samples taken from the same individual within 7 days of each other were compared using different masking strategies. The top panel describes the published, deployed method of masking. In the panels below, genes in the regions shown in Figure 5B are additionally masked (i.e. ignored from pairwise comparisons).

## DISCUSSION

Here, we describe an approach which we term adaptive masking. This involves monitoring the minor variant frequency across a bacterial genome to which sequencing reads have been mapped; as such, it measures an end product of next generation sequencing processes, taking into account the natural sequence variation in the samples studied, as well as the impact of DNA extraction and library construction technologies, and the performance of the mapping and filtering software used.

Using this approach, we demonstrated a significant positive association between amounts of non-Mycobacterial bacterial DNA and minor variant frequencies in a subset of the mapped genome: significant increases of more than 1.5 fold were observed in 242/8006 regions examined, which together cover about 5% of the *M. tuberculosis* genome. Among these 242 regions we identified four ‘high variation’ regions in which minor variant frequencies are very strongly influenced by non-Mycobacterial bacterial DNA quantities, with fold-increases in minor variant frequencies of >5 in the presence of >20% non-Mycobacterial bacterial DNA. Importantly, if non-Mycobacterial bacterial DNA concentrations are low (<1% of bacterial DNA present), as occurred in retrospective studies when Mycobacteria were subcultured on Lowenstein-Jensen slopes prior to sequencing, increased variation is not observed in these regions.

The exclusion of the four ‘high variation’ regions from base calling by a clinically deployed *M. tuberculosis* pipeline markedly reduced reported variation between samples derived from the same patient in a short time period. In particular, prior to exclusion of the four high variation regions, in a test set derived from 234 individuals, 5.2% of intra-patient pairs examined differed by 5 SNV or more, with the majority of SNV differences observed in these pairs being >20. Multiple studies indicate this is biologically implausible [3, 5, 7], and after exclusion of the four ‘high variation’ regions comprising only 0.2% of the genome, no pairs had variation of 5 SNV or more. This suggests that using standard ‘masking’ and DNA extraction from liquid media false positive variation is reported in a small number of sites in a non-Mycobacterial bacterial DNA dependent manner. Put alternatively, non-Mycobacterial bacterial DNA acts as an interfering substance [20] for relatedness measurements.

A potential limitation of this work is that this approach studies pre-specified regions of the genome, specifically coding regions and intergenic regions. This approach was chosen to avoid the challenges of analysing the 4.4 x 10^6^ bases of the *M. tuberculosis* genome individually, with concomitant loss of statistical power. Therefore, as described the method may neither detect, nor allow selective masking of, small regions with high minor variant frequencies within genes. A second limitation is that we did not explore the use of metagenomics classifiers, such as Kraken, to identify non-Mycobacterial ‘interfering’ DNA and eliminate it prior to mapping to the *M. tuberculosis* genome. We chose not to do this because we observed that for the highly conserved *rrs* genes, metagenomic classifiers cannot confidently assign reads to a genus level, likely because there is insufficient sequence variation within *rrs* to allow this. Despite these limitations, the strategy chosen appears to be of use clinically, based on the reduction in likely false positive variation between serial samples from individuals.

The clinical of use of next generation sequencing is rapidly increasing [1, 4, 22]. Clinically deployed bioinformatic pathways reporting microbial identity, resistotyping and relatedness information are complex, multistep processes whose outputs are dependent on specimen decolonisation, selective culture, DNA extraction, library construction, DNA sequencing, and bioinformatic analysis [4].

Reagent batches, software versions, and equipment involved in the process are all subject to change over time. Generalizable to other organisms and mapping pipelines, the adaptive masking approach we described here will have application in monitoring the performance of such processes quantitatively and preventing the calling of false positive variation in the context of clinically deployed genomics. Additionally, it offers a route to monitoring the performance of critical steps in the laboratory and bioinformatics processes determining microbial relatedness in the context of a closed loop, auditable process as required by standard laboratory accreditation systems, such as ISO15189.

## ACKNOWLEDGEMENTS

This study is supported by the Health Innovation Challenge Fund (a parallel funding partnership between the Wellcome Trust [WT098615/Z/12/Z] and the Department of Health [grant HICF-T5-358]) and NIHR Oxford Biomedical Research Centre. Professor Derrick Crook is affiliated to the National Institute for Health Research Health Protection Research Unit (NIHR HPRU) in Healthcare Associated Infections and Antimicrobial Resistance at University of Oxford in partnership with Public Health England. Professor Crook is based at University of Oxford. The views expressed are those of the author(s) and not necessarily those of the NHS, the NIHR, the Department of Health or Public Health England. The sponsors of the study had no role in study design, data collection, data analysis, data interpretation, or writing of the report. The corresponding author had full access to all the data in the study and had final responsibility for the decision to submit for publication.

## AVAILABILITY OF DATA

Sample meta data and excel versions of the supplementary data is available here: https://ora.ox.ac.uk/objects/uuid:88d93ec1-2757-4e46-83f4-dbdfc01b5343.

## FIGURES AND FIGURE LEGENDS

**Figure S1.**
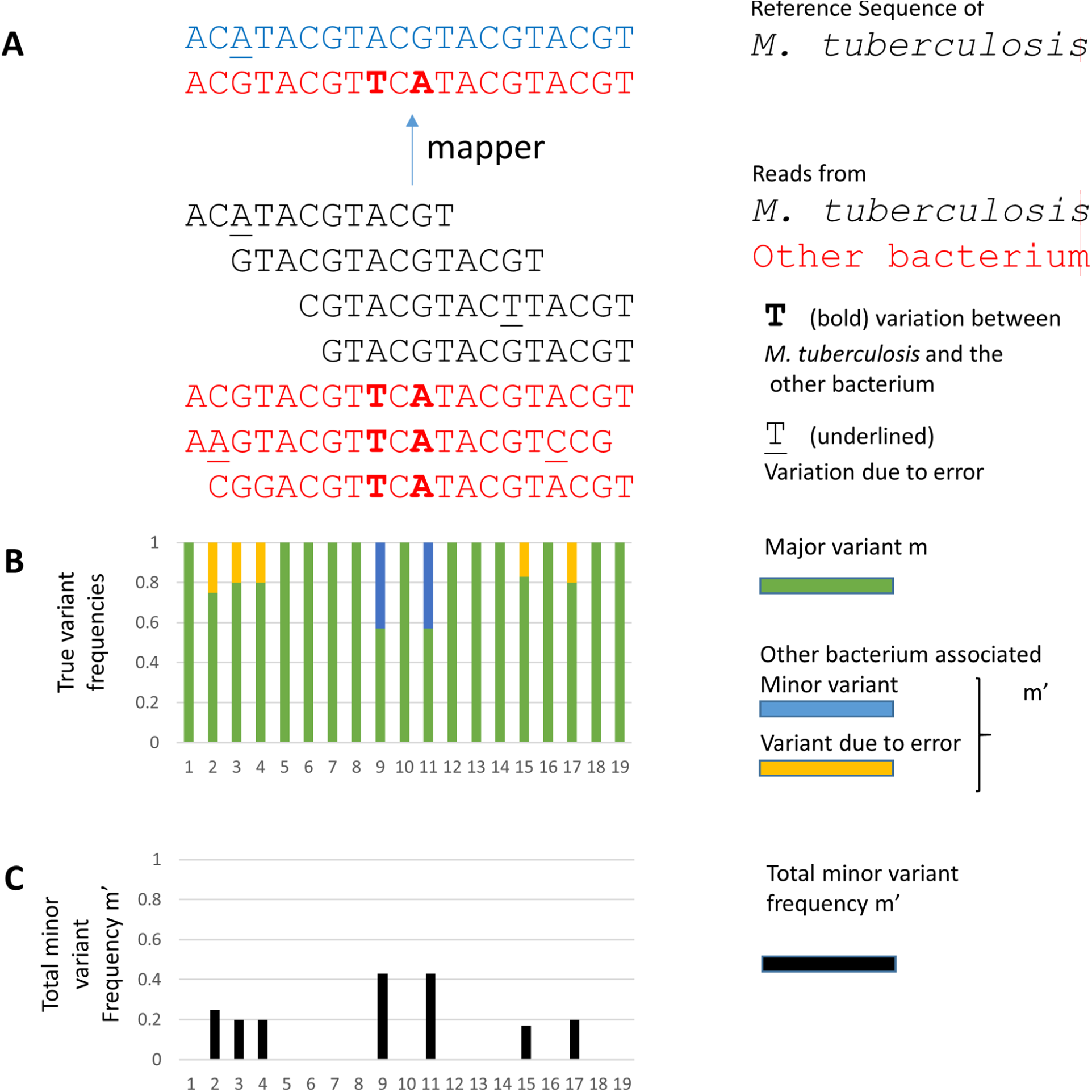
Illustration of minor variant frequencies. Legend: (A) Short read sequencing data, either from *M. tuberculosis* or from other bacteria (red) is mapped to a reference gene. Minor variants (B) can result either from sequencing error (underlined), or due to alignment of non-*Mycobacterium tuberculosis* DNA to the reference. Variation of both types contribute to the minor variant frequency (C).

**Figure S2.**
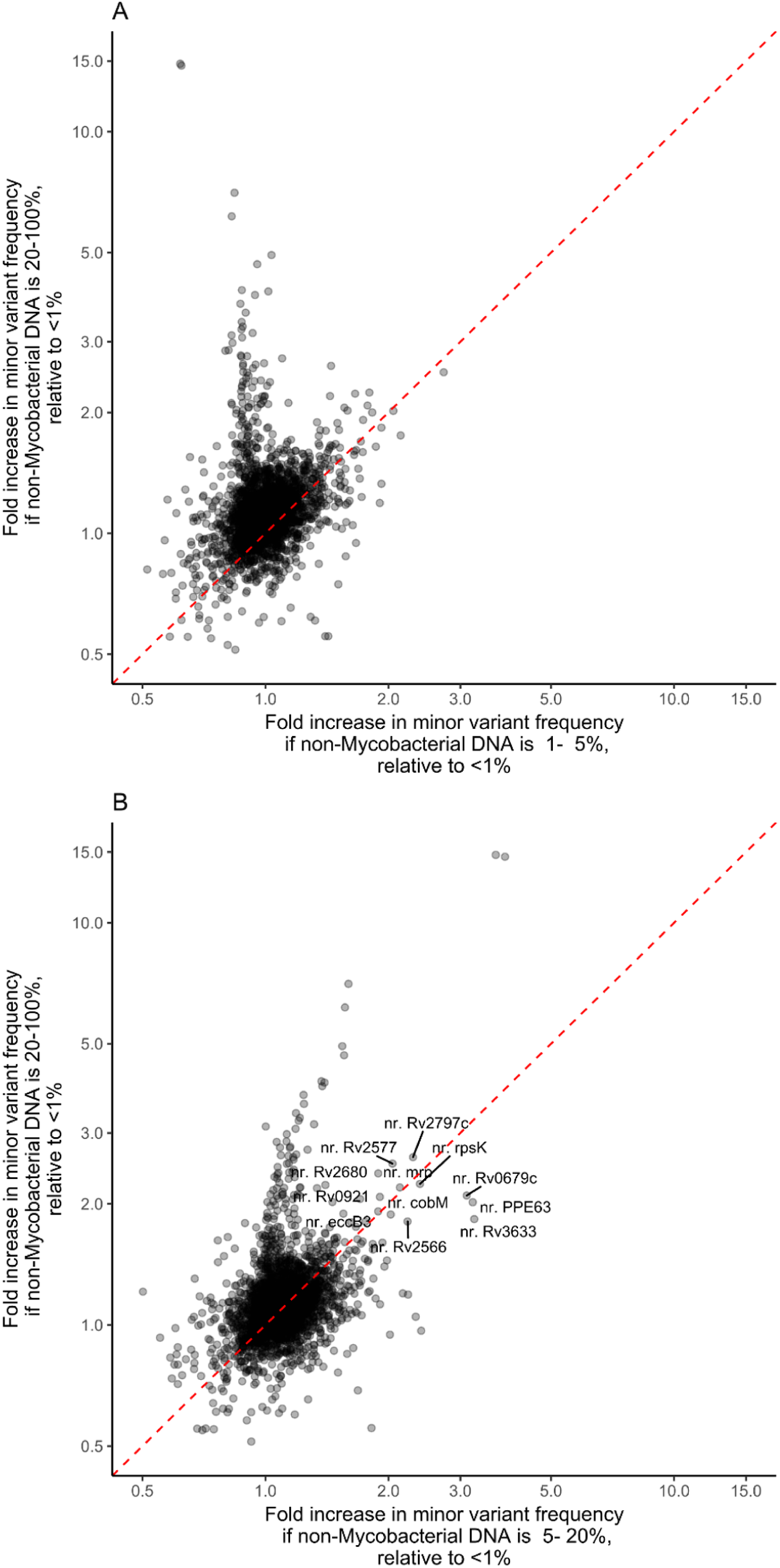
Minor variant frequencies with increasing non-Mycobacterial bacterial DNA. Legend: Estimated rate ratios derived from Poisson modelling of minor variant frequencies relative to frequencies when &1% non-Mycobacterial bacterial DNA was present. (A) compares rate ratios with 1-5% vs. &20% non-Mycobacterial bacterial DNA; (B) compares 5-20% vs. &20% non-Mycobacterial bacterial DNA. In (B), regions with similar rate ratios when 5-20% and &20% non-Mycobacterial bacterial DNA are annotated.

**Figure S3.**
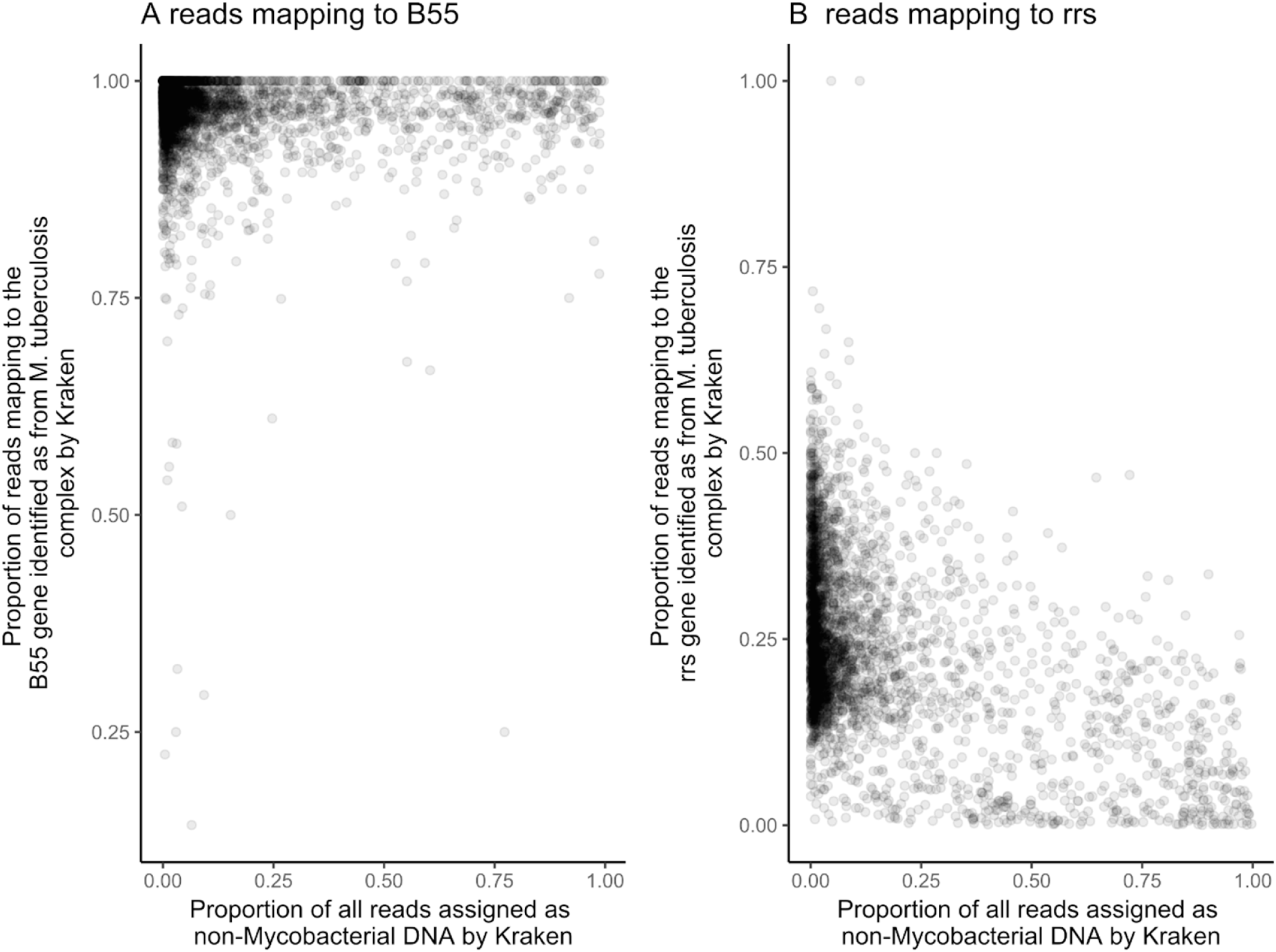
Identification of reads in selected regions by Kraken. Legend: The proportion of reads mapped to (A) the B55 gene (B) the rrs genes which are identified as belonging to the *M. tuberculosis* complex by Kraken are shown, as is their relationship with the estimate of the amount of non-Mycobacterial bacterial DNA in the all reads.

## REFERENCES

1. De Silva D, Peters J, Cole K, Cole MJ, Cresswell F, Dean G, et al. Whole-genome sequencing to determine transmission of Neisseria gonorrhoeae: an observational study. The Lancet Infectious diseases. 2016;16(11):1295–303. Epub 2016/10/30. doi:10.1016/s1473-3099(16)30157-8. PubMed PMID: 27427203; PubMed Central PMCID: PMCPMC5086424.

2. Gordon NC, Pichon B, Golubchik T, Wilson DJ, Paul J, Blanc DS, et al. Whole-Genome Sequencing Reveals the Contribution of Long-Term Carriers in Staphylococcus aureus Outbreak Investigation. Journal of clinical microbiology. 2017;55(7):2188–97. Epub 2017/05/05. doi:10.1128/jcm.00363-17. PubMed PMID: 28468851; PubMed Central PMCID: PMCPMC5483921.

3. Walker TM, Lalor MK, Broda A, Ortega LS, Morgan M, Parker L, et al. Assessment of Mycobacterium tuberculosis transmission in Oxfordshire, UK, 2007-12, with whole pathogen genome sequences: an observational study. The Lancet Respiratory medicine. 2014;2(4):285–92. Epub 2014/04/11. doi:10.1016/s2213-2600(14)70027-x. PubMed PMID: 24717625; PubMed Central PMCID: PMCPMC4571080.

4. Pankhurst LJ, Del Ojo Elias C, Votintseva AA, Walker TM, Cole K, Davies J, et al. Rapid, comprehensive, and affordable mycobacterial diagnosis with whole-genome sequencing: a prospective study. The Lancet Respiratory medicine. 2016;4(1):49–58. Epub 2015/12/17. doi:10.1016/s2213-2600(15)00466-x. PubMed PMID: 26669893; PubMed Central PMCID: PMCPMC4698465.

5. Bryant JM, Harris SR, Parkhill J, Dawson R, Diacon AH, van Helden P, et al. Whole-genome sequencing to establish relapse or re-infection with Mycobacterium tuberculosis: a retrospective observational study. The Lancet Respiratory medicine. 2013;1(10):786–92. Epub 2014/01/28. doi:10.1016/s2213-2600(13)70231-5. PubMed PMID: 24461758; PubMed Central PMCID: PMCPMC3861685.

6. Casali N, Nikolayevskyy V, Balabanova Y, Harris SR, Ignatyeva O, Kontsevaya I, et al. Evolution and transmission of drug-resistant tuberculosis in a Russian population. Nature genetics. 2014;46(3):279–86. Epub 2014/01/28. doi:10.1038/ng.2878. PubMed PMID: 24464101; PubMed Central PMCID: PMCPMC3939361.

7. Guerra-Assuncao JA, Crampin AC, Houben RM, Mzembe T, Mallard K, Coll F, et al. Large-scale whole genome sequencing of M. tuberculosis provides insights into transmission in a high prevalence area. eLife. 2015;4. Epub 2015/03/04. doi:10.7554/eLife.05166. PubMed PMID: 25732036; PubMed Central PMCID: PMCPMC4384740.

8. Guerra-Assuncao JA, Houben RM, Crampin AC, Mzembe T, Mallard K, Coll F, et al.Recurrence due to relapse or reinfection with Mycobacterium tuberculosis: a whole-genome sequencing approach in a large, population-based cohort with a high HIV infection prevalence and active follow-up. The Journal of infectious diseases. 2015;211(7):1154–63. Epub 2014/10/23. doi:10.1093/infdis/jiu574. PubMed PMID: 25336729; PubMed Central PMCID: PMCPMC4354982.

9. Morgulis A, Gertz EM, Schaffer AA, Agarwala R., A fast and symmetric DUST implementation to mask low-complexity DNA sequences. Journal of computational biology: a journal of computational molecular cell biology. 2006;13(5):1028–40. Epub 2006/06/27. doi:10.1089/cmb.2006.13.1028. PubMed PMID: 16796549.

10. Bedell JA, Korf I, Gish W. MaskerAid: a performance enhancement to RepeatMasker. Bioinformatics (Oxford, England). 2000;16(11):1040–1. Epub 2001/02/13. PubMed PMID: 11159316.

11. Langmead B, Trapnell C, Pop M, Salzberg SL., Ultrafast and memory-efficient alignment of short DNA sequences to the human genome. Genome biology. 2009;10(3):R25. Epub 2009/03/06. doi:10.1186/gb-2009-10-3-r25. PubMed PMID: 19261174; PubMed Central PMCID:PMCPMC2690996.

12. Li H, Ruan J, Durbin R., Mapping short DNA sequencing reads and calling variants using mapping quality scores. Genome research. 2008;18(11):1851–8. Epub 2008/08/21. doi:10.1101/gr.078212.108. PubMed PMID: 18714091; PubMed Central PMCID: PMCPMC2577856.

13. Li H, Durbin R., Fast and accurate short read alignment with Burrows-Wheeler transform. Bioinformatics (Oxford, England). 2009;25(14):1754–60. Epub 2009/05/20. doi:10.1093/bioinformatics/btp324. PubMed PMID: 19451168; PubMed Central PMCID: PMCPMC2705234.

14. Lunter G, Goodson M.Stampy: a statistical algorithm for sensitive and fast mapping of Illumina sequence reads. Genome research. 2011;21(6):936–9. Epub 2010/10/29. doi:10.1101/gr.111120.110. PubMed PMID: 20980556; PubMed Central PMCID: PMCPMC3106326.

15. Vijay S, Dalela G., Prevalence of LRTI, in Patients Presenting with Productive Cough and Their Antibiotic Resistance Pattern. Journal of clinical and diagnostic research: JCDR. 2016;10(1):Dc09–12. Epub 2016/02/20. doi:10.7860/jcdr/2016/17855.7082. PubMed PMID: 26894065; PubMed Central PMCID: PMCPMC4740592.

16. SMI B 40: Investigation of specimens for Mycobacterium species. London: Public Health England, 2017.

17. Quan TP, Bawa Z, Foster D, Walker T, Del Ojo Elias C, Rathod P, et al. Evaluation of whole genome sequencing for Mycobacterial species identification and drug susceptibility testing in a clinical setting: a large-scale prospective assessment of performance against line-probe assays and phenotyping. Journal of clinical microbiology. 2017. Epub 2017/11/24. doi:10.1128/jcm.01480-17. PubMed PMID: 29167290.

18. Comas I, Coscolla M, Luo T, Borrell S, Holt KE, Kato-Maeda M, et al. Out-of-Africa migration and Neolithic coexpansion of Mycobacterium tuberculosis with modern humans. Nature genetics. 2013;45(10):1176–82. Epub 2013/09/03. doi:10.1038/ng.2744. PubMed PMID: 23995134; PubMed Central PMCID: PMCPMC3800747.

19. Hatherell HA, Colijn C, Stagg HR, Jackson C, Winter JR, Abubakar I. Interpreting whole genome sequencing for investigating tuberculosis transmission: a systematic review. BMC medicine. 2016;14:21. Epub 2016/03/24. doi:10.1186/s12916-016-0566-x. PubMed PMID: 27005433; PubMed Central PMCID: PMCPMC4804562.

20. Standardisation IOf. Medical Laboratories - Requirements for Quality and Competence. 2012.

21. Nasidze I, Li J, Quinque D, Tang K, Stoneking M. Global diversity in the human salivary microbiome. Genome research. 2009;19(4):636–43. doi:10.1101/gr.084616.108. PubMed PMID: PMC2665782.

22. Bradley P, Gordon NC, Walker TM, Dunn L, Heys S, Huang B, et al. Rapid antibiotic-resistance predictions from genome sequence data for Staphylococcus aureus and Mycobacterium tuberculosis. Nature communications. 2015;6:10063. Epub 2015/12/22. doi:10.1038/ncomms10063. PubMed PMID: 26686880; PubMed Central PMCID: PMCPMC4703848.

23. Wood DE, Salzberg SL. Kraken: ultrafast metagenomic sequence classification using exact alignments. Genome biology. 2014;15(3):R46. Epub 2014/03/04. doi:10.1186/gb-2014-15-3-r46. PubMed PMID: 24580807; PubMed Central PMCID: PMCPMC4053813.

24. Street TL, Sanderson ND, Atkins BL, Brent AJ, Cole K, Foster D, et al. Molecular Diagnosis of Orthopedic-Device-Related Infection Directly from Sonication Fluid by Metagenomic Sequencing. Journal of clinical microbiology. 2017;55(8):2334–47. Epub 2017/05/12. doi:10.1128/jcm.00462-17. PubMed PMID: 28490492; PubMed Central PMCID: PMCPMC5527411.

25. Li H, Handsaker B, Wysoker A, Fennell T, Ruan J, Homer N, et al. The Sequence Alignment/Map format and SAMtools. Bioinformatics (Oxford, England). 2009;25(16):2078–9. Epub 2009/06/10. doi:10.1093/bioinformatics/btp352. PubMed PMID: 19505943; PubMed Central PMCID: PMCPMC2723002.

26. Walker TM, Kohl TA, Omar SV, Hedge J, Del Ojo Elias C, Bradley P, et al. Whole-genome sequencing for prediction of Mycobacterium tuberculosis drug susceptibility and resistance: a retrospective cohort study. The Lancet Infectious diseases. 2015;15(10):1193–202. Epub 2015/06/28. doi:10.1016/s1473-3099(15)00062-6. PubMed PMID: 26116186; PubMed Central PMCID: PMCPMC4579482.

27. Mazariegos-Canellas O, Do T, Peto T, Eyre DW, Underwood A, Crook D, et al. BugMat and FindNeighbour: command line and server applications for investigating bacterial relatedness. BMC bioinformatics. 2017;18(1):477. Epub 2017/11/15. doi:10.1186/s12859-017-1907-2. PubMed PMID: 29132318; PubMed Central PMCID: PMCPMC5683244.

28. Coll F, McNerney R, Guerra-Assuncao JA, Glynn JR, Perdigao J, Viveiros M, et al. A robust SNP barcode for typing Mycobacterium tuberculosis complex strains. Nature communications. 2014;5:4812. Epub 2014/09/02. doi:10.1038/ncomms5812. PubMed PMID: 25176035; PubMed Central PMCID: PMCPMC4166679.

